# DBERlibR: Automated Data Analysis for Discipline-Based Education Research

**DOI:** 10.1101/2022.08.24.504993

**Authors:** Changsoo Song, Resa Helikar, Wendy M. Smith, Tomáš Helikar

## Abstract

Discipline-Based Education Research (DBER) scientists repeatedly analyze assessment data to ensure question items’ reliability and examine the efficacy of a new educational intervention. Analyzing assessment data comprises multiple steps and statistical techniques that consume much of researchers’ time and are error-prone. While education research continues to grow across many disciplines of science, technology, engineering, and mathematics (STEM), the DBER community lacks tools to streamline education research data analysis. DBERlibR—an R package to streamline and automate DBER data processing and analysis—fills this gap. The package reads user-provided assessment data, cleans them, merges multiple datasets (as necessary), checks assumption(s) for specific statistical techniques (as necessary), applies various statistical tests (e.g., one-way analysis of covariance, one-way repeated-measures analysis of variance), and presents and interprets the results all at once. By providing the most frequently used analytic techniques, this package will contribute to DBER by facilitating the creation and widespread use of evidence-based knowledge and practices. The outputs contain a sample interpretation of the results for users’ convenience. User inputs are minimal; they only need to prepare the data files as instructed and type a function in RStudio to conduct a specific data analysis.

## 1. Introduction

Discipline-based education research (DBER) aims to improve educational practices in the science, technology, engineering, and mathematics (STEM) disciplines, including physics, life sciences, engineering, and mathematics (National Research Council 2012; Singer and Smith 2013). DBER scientists implement a range of educational interventions/methods that are deeply grounded in a particular discipline and pursue evidence-based knowledge and practices to enhance teaching and learning in STEM disciplines (Colaninno 2019; Henderson, Connolly, Dolan, Finkelstein, Franklin, Malcom, Rasmussen, Redd, and John 2017). The pursuit of evidence-based practices entails continuous development and testing of various learning interventions and repeated collection, cleaning, integration, and analysis of educational assessment data.

However, cleaning data (e.g., treating missing values, handling incorrect values and outliers) is a daunting task for education researchers, as is performing statistical data analyses (e.g., item analysis, repeated measures analysis of variance). Erroneous coding and inconsistent data generation impede data cleaning. Researchers must go through each question item in the dataset and look for potential errors, a process that can be very time-consuming and error-prone (Petersen and Ekstrøm 2019). Depending on the evaluation design, different statistical techniques are employed to analyze the assessment data; investigators merge the data, test assumptions required for parametric techniques, or employ non-parametric techniques as necessary (Naskar and Das 2018), which are also time-consuming and error-prone. Researchers usually perform these tasks individually using multiple software packages and functions because of the current piecemeal statistical data analysis tools. No single tool brings the whole pipeline together for a more streamlined assessment data analysis for DBER, making it challenging to increase overall study reproducibility.

Here, we present DBERlibR, an easy-to-use R package to streamline and automate DBER data processing and analysis, helping investigators to avoid input errors, increase analysis speed, and improve the consistency of result output.

## 2. Assessment Data Preparation

The minimum requirement to use installed DBERlibR is a user-provided assessment data file (ADF) that contains graded assessment data for all study subjects. The ADF should include only subject identifiers and assessment items (Figure 1). No other data like demographic information should be included in the ADF. Users should save other data in a separate file along with the identifiers - see the demographic data format below. The assessment data should be binary (i.e., 1 for correct answers, 0 for incorrect answers). The ADF can have missing values (these are treated by DBERlibR automatically). The ADF should be in a “CSV” (comma separated values) format.

**Figure 1:**
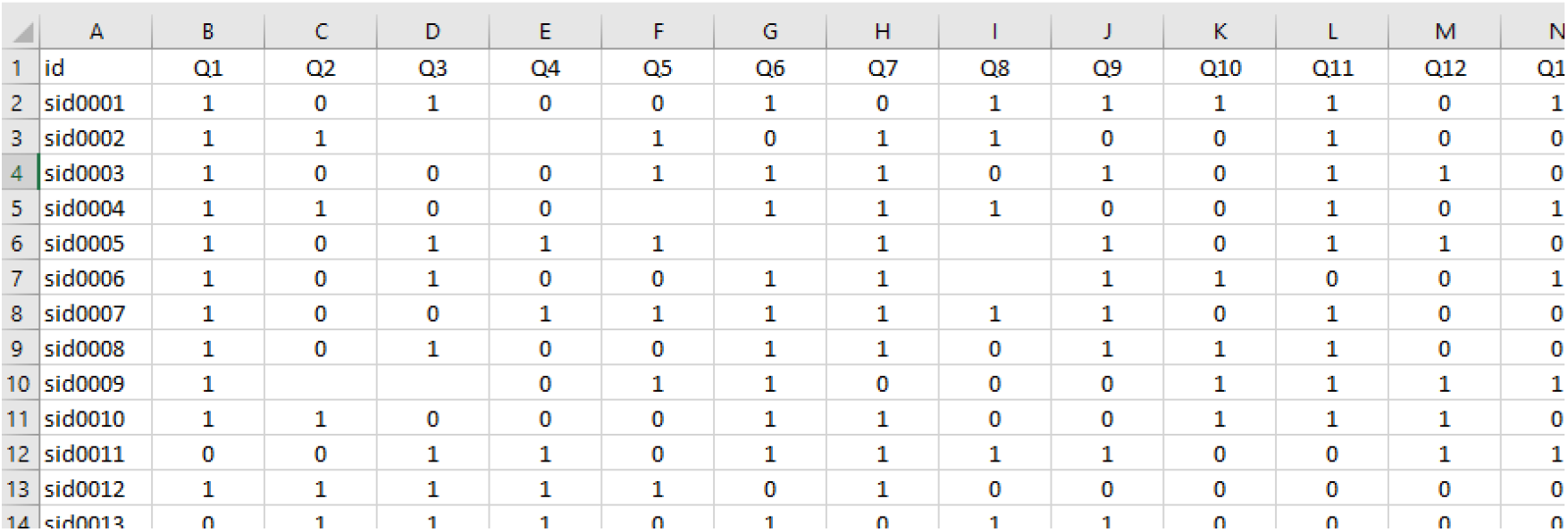
Example of an Assessment Data File Format.

**Figure 2:**
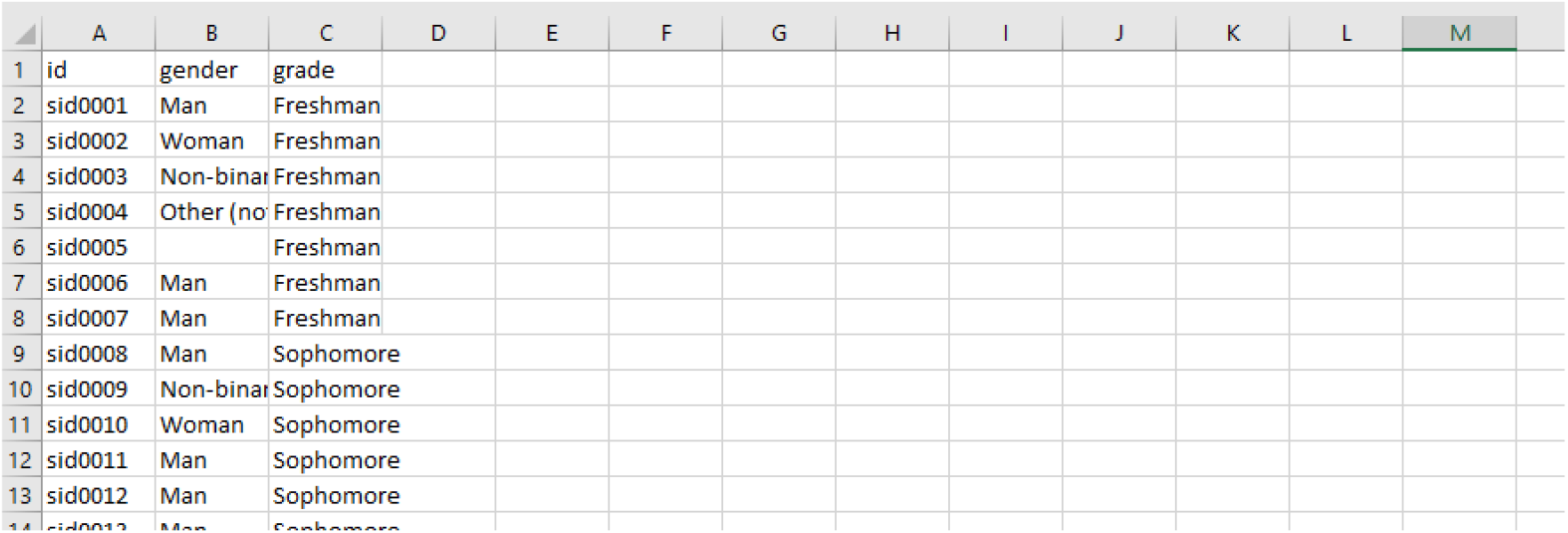
Demographic Data Format

Figure 1 shows an ADF file example in which the assessment items were named Q1-n. Users can name the assessment items freely. Blank cells indicate missing values (e.g., skipped answers). These blank cells will be treated as incorrect in the data preparation process (per standard practices). Users must include only one header row.

DBERlibR supports using and analyzing multiple ADFs to support, for example, control and treatment groups or pre-treatment and post-treatment assessments. The Users need to save the data file exactly the same as the sample data files (use the same file names):

- “data_treat_pre.csv” for the treatment group’s pre-test data;
- “data_treat_post.csv” for the treatment group’s post-test data;
- “data_treat_post2.csv” for the treatment group’s second post-test data;
- “data_ctrl_pre.csv” for the control group’s pre-test data;
- “data_ctrl_post.csv” for the control group’s post-test data; and
- “data_ctrl_post2.csv” for the control group’s second post-test data.

Data file(s) should be saved in a folder set as a working directory from which the functions in this package import data file(s).

If users have students’ demographic information to examine the differences across various subgroups (e.g., by gender, ethnicity, or year in school), these data can be provided to DBERlibR in a separate file along with identifiers (the same identifiers as in the ADF(s)).

## 3. Overview of DBERlibR Features

DBERlibR automatically reads user-provided assessment data, cleans inaccurate or missing values, merges multiple datasets (as necessary), checks assumption(s) for a specific statistical technique (as necessary), runs the main assessment data analysis, and presents and assists with data interpretation in a streamlined fashion. The current version of DBERlibR covers item analysis, paired-samples t-test (and Wilcoxon signed-rank test as necessary), independent samples t-test (and Mann-Whitney U test as necessary), one-way analysis of covariance (ANCOVA), one-way repeated measures analysis of variance (ANOVA) (and Friedman test as necessary), two-way repeated-measures ANOVA, and one-way analysis of variance (and Kruskal-Wallis rank-sum test as necessary).

If users’ data contain multiple-choice answers, they must transform them into a binary format of 1 for correct answers and 0 for incorrect answers. The function of **multi_to_binary(csv_data)** helps users do this job automatically. Before running this function, users need to create and save a CSV file with answer keys for all questions in the data file.

The function of **item_analysis(csv_data)** provides item difficulty and item discrimination scores. The item difficulty refers to the proportion of students correctly answering the question item, and the item discrimination refers to the relationship between how well students did on the item and their total test performance. Users can utilize this function to improve the accuracy of their question items in assessing student performance. Outputs from this function include a graph with ordered scores (low to high) so that users can instantly identify problematic question items (refer to the section of Case Study for more details).

The function of **paired_samples()** helps users compare pre-test and post-test scores. The paired samples t-test requires an assumption of normality to be satisfied. Therefore, this function automatically conducts the Shapiro-Wilk normality test, which tests the null hypothesis that a sample (*x*_*1*,…,_*x*_*n*_) came from a normally distributed population. The test statistic (*W*) is calculated by:

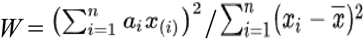

where x_(i)_ is the i^th^-smallest number in the sample and 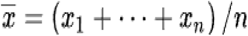 is the sample mean (Shapiro and Wilk 1965). If the assumption of normality is satisfied, the function runs the parametric paired samples t-test.

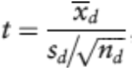

where *t* is the test statistic, *<u>x</u>*_*d*_ is the sample mean difference score, *s*_*d*_ is the standard deviation of the sample difference scores, and *n*_*d*_ is the number of paired observations in the sample (Stone 2010).

If the normality assumption is not satisfied, the function runs the non-parametric Wilcoxon signed-rank test.

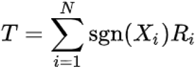

where *T* is the test statistic, sgn is the sign function:sgn(*x*) = 1 if *x* > 0 and sgn(*x*) = -1 if *x* < 0, and *R*_*i*_ is the number of *j* for which |*X*_*j*_|≤|*X*_*i*_| (Siegel 1956; Wilcoxon 1945). Then, the function provides results and a brief interpretation of the results.

The function of **independent_samples()** compares two independent groups (e.g., intervention vs. control group). The independent samples t-test requires assumptions of normality and equal variances, so this function automatically checks the assumptions. If the assumption of normality is satisfied, the function runs the parametric independent samples t-test:

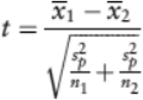

where *t* is the test statistic, *<u>x</u>*_*1*_ is the sample mean of group 1, *<u>x</u>*_*2*_ is the sample mean of group 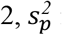 is the pooled estimate of variance, and *n*_*1*_ is the sample size of group 1, *n*_*2*_ is the sample size of group 2 (Stone 2010). If the assumption of normality is not satisfied, the function runs the non-parametric Mann-Whitney U test:

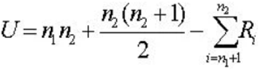

where U is Mann-Whitney U test, *n*_*1*_ is the sample size of group 1, *n*_*2*_ is the sample size of group 2, and *R*_*i*_ is the rank of the sample size (Hinton 2010). Then, the function provides either result with or without equal the variances assumption (in case of the parametric test).

The function **one_way_repeated_anova()** analyzes the changes over time (e.g., the difference among pre-, post-, and post2-test scores of the intervention group). The one-way repeated measures ANOVA requires the assumptions of normality, equal variances, and sphericity (refer to Acheson 2010 for details) to be satisfied, so this test begins with the normality test. If the assumption of normality is satisfied, the function runs runs the parametric one-way repeated ANOVA (refer to Gueorguieva and Krystal 2004 for details) If the normality assumption is not satisfied, the function runs the non-parametric Friedman test:

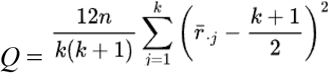

where *Q* is the test statistic, *n* is the number of rows (blocks), *k* is the number of columns (treatments), and 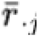:

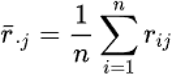

where r_ij_ is the rank of ***x***_***ij***_ within block *i* (Friedman 1937). Then, the function provides results either with or without equal the variances assumption.

Meanwhile, the function **two_way_repeated_anova()** analyzes the interaction effect between time (pre-post-post2) and conditions for the intervention group. The function automatically checks the assumptions of normality and sphericity that need to be satisfied. If there is a significant difference between times or conditions, the function automatically conducts post hoc analyses (refer to Acheson 2010 for details).

The function **one_way_ancova()** analyzes the difference between two groups (e.g., intervention vs. control group) with a covariate (e.g., pre-test scores) controlled. The function automatically checks the assumptions of linearity, normality, equal variance, and homogeneity of regression line slopes that need to be satisfied and then conducts one-way ANCOVA:

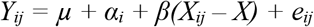

where *Y*_*ij*_ is the outcome for person *i* in group *j* (e.g., *j*=1 for control, *j*=2 for intervention), *µ* is the grand mean of *Y, α*_*i*_ is the effect of intervention j, *β* is the slope of the regression line, *X*_*ij*_ is the covariate value for person i in group j, and *e*_*ij*_ is a normally distributed residual or error term with a mean of zero and a variance 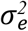 (van Breukelen 2010). Then, the function runs post hoc analyses to report, depending on the test result.

Finally, the function **demo_group_diff(csv_data)** helps users analyze the difference between sub-groups of a demographic variable (e.g., gender, grade). The function automatically checks the assumptions of normality and equal variances. If the assumption of normality is satisfied, the function runs the parametric one-way ANOVA:

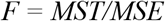

where *F* is the test statistic, *MST* is the mean square for treatment, and *MSE* is the mean square error (Wahed and Tang 2010). If the normality is not satisfied, the function runs the non-parametric Mann-Whitney U test (if two sub-groups) or Kruskal-Wallis test (if three or more sub-groups. The Kruskal-Wallis test is run by:

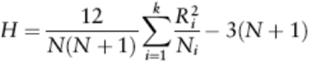

where *H* is the test statistic, *N*_*i*_ is the number of all observations by *N*, and *R*_*i*_ is the sum of ranks for all *k* samples (from *i* = 1 to *i* = *k*) (Schmidt 2010).

All functions automatically clean data and merge different datasets as necessary (e.g., merge pre-test and post-test groups, bind intervention and control groups) before analyzing data. They generate a summary statement and interpretation of the results to report. Figure 3 graphically illustrates the procedures of data analysis by DBERlibR.

**Figure 3:**
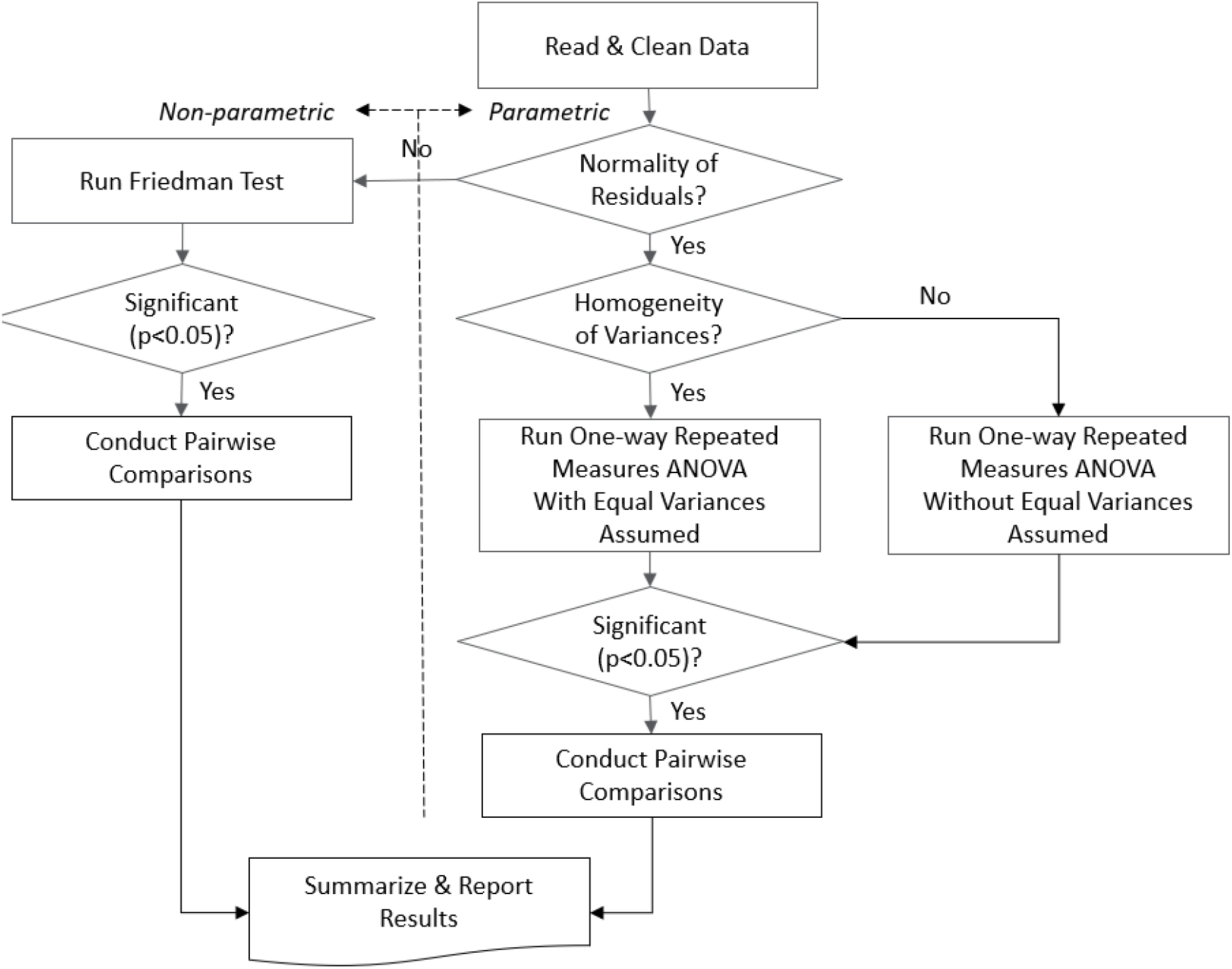
DBERlibR Workflow - An Illustration for the One-way Repeated Measures ANOVA

The following section illustrates using three of DBERlibR’s functions: **item_analysis, one-way_ancova**, and **one_way_repeated_anova**, to demonstrate what DBERlibR offers to help users with data analysis.

## 4. Case Study: Impact of hands-on systems modeling on learning in biochemistry

This section illustrates and exemplifies how DBERlibR can be used in a specific discipline-based education research study. The presented case study is based on our recent work that investigated whether interactive computer simulations of metabolic networks can increase students’ ability to recognize how individual interactions between biochemical components affect the behavior of a system under different conditions (Booth, Song, Howell, Rasquinha, Saska, Helikar, Sikich, Couch, van Dijk, Roston, and Helikar 2021). Data for this study were collected from two upper-level college Biochemistry class sections to conduct a DBER project to answer a research question: “Is computational modeling effective in enhancing students’ understanding of complex biological processes?” A non-equivalent group design was employed to collect and analyze data. One section of the class was the intervention group, whereas the other section served as control. The data were collected from both sections before and after deploying a computational modeling lesson in the intervention group. The number of students from both sections was 87 (40 from the intervention group and 47 from the control group). The research team developed a set of question items for the assessment and conducted a series of item analyses to evaluate the level of difficulty and discrimination of individual question items to secure the reliability and accuracy of the assessment. DBERlibR can help develop an appropriate assessment instrument through item analysis and answer the research question through one-way ANCOVA.

### 4.1. Item analysis: “item_analysis(csv_data)”

The item analysis function requires users to type an ADF name, as shown in the sample code below. Users can input any of the data file names (i.e., “data_treat_pre.csv,” “data_treat_post.csv,” “data_ctrl_pre.csv,” “data_ctrl_post.csv,” “data_treat_post2.csv” and data_ctrl_post2.csv”) as they become available for item analysis.

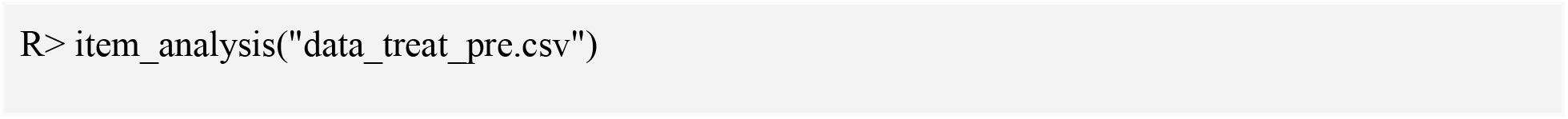

Then, the function automatically reads and cleans the data (e.g., treating wrong inputs/typos as missing, converting missing values to “0”). The function returns a warning that “Skipped answers will be treated as incorrect in this package, and too many skipped answers may skew the results of data analysis.” Users can define too many skipped answers with a percentage, and students with too many skipped answers will be excluded from data analysis to prevent skewed results. For instance, if users input 15%, students who skipped more than 15% of the question will be deleted.

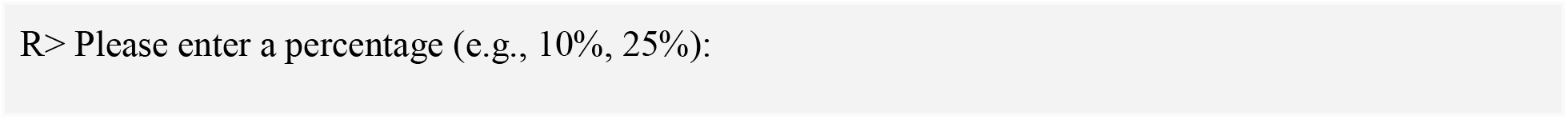

After automatic data cleaning, the function calculates difficulty and discrimination scores, displays the results in RStudio, and exports the results to an Excel file, which is saved in the working directory designated by users. The Excel file name follows the data file name used for item analysis. For example, if users have called “data_treat_pre.csv” in the function, then the output Excel file name contains “treat_pre.” The function also generates plots in the PNG file format to visualize the results (Figures 4 and 5) so that users (teachers/instructors) can instantly identify question items presenting room for improvement. The PNG file name also follows the rule as the data file name, so users can easily find the item analysis results in the working directory.

**Figure 4:**
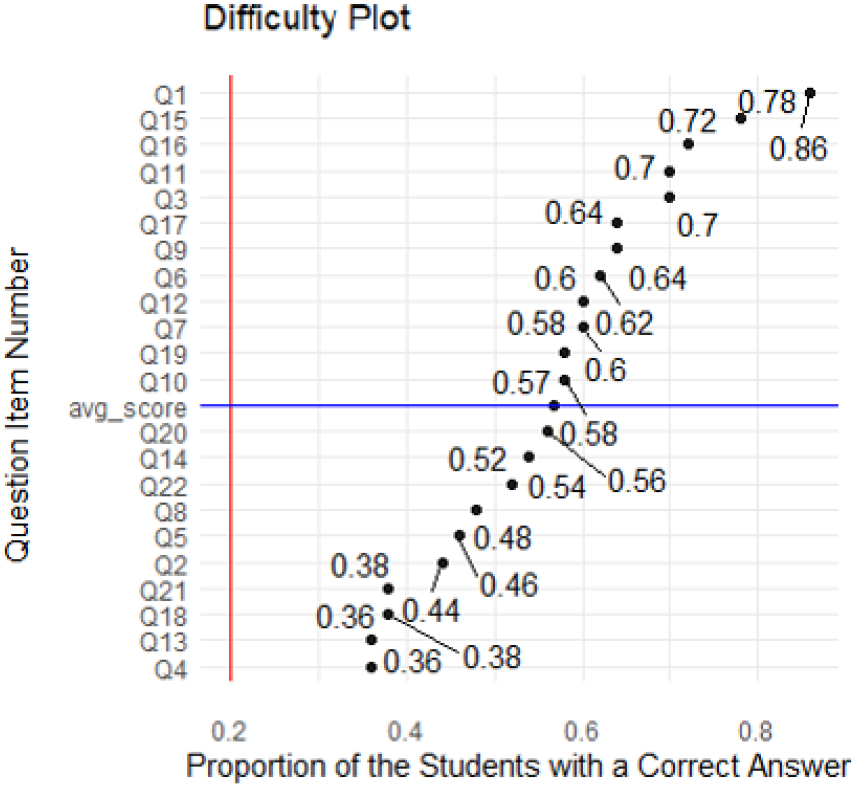
Item Difficulty Analysis

**Figure 5:**
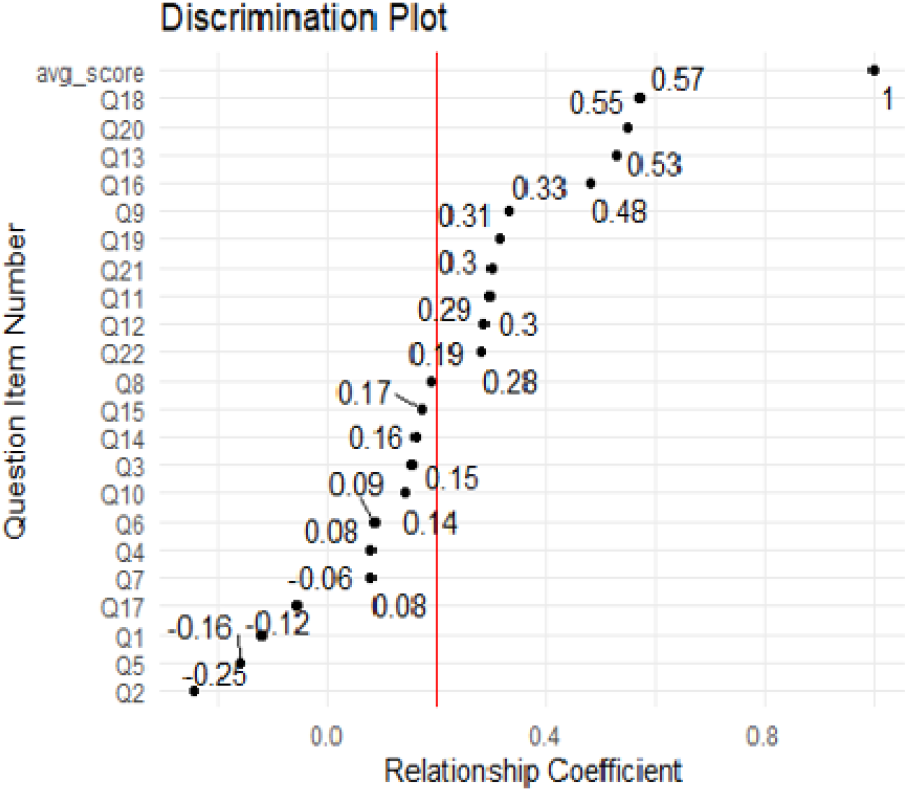
Item Discrimination Analysis

Item difficulty refers to the proportion of students that correctly answered the question item. These proportions are ordered by size in the difficulty plot (Figure 4). Question items for which less than 20% of students (0.2) got the correct answer are considered “too difficult.” These questions may not assess students’ knowledge or skills accurately. The vertical red-colored line represents the threshold so that users can instantly find the question items that warrant their attention for improvement (e.g., eliminate or rephrase/reword). The difficulty plot above does not feature any item whose difficulty is lower than 0.2.

Item discrimination indicates the relationship between the performance of students on a given item and their total test performance. The relationship coefficients are ordered by size in the discrimination plot (Figure 5). A coefficient lower than 0.2 reflects a lack of discrimination by the item. The vertical red-colored line represents the threshold so that users can instantly identify potentially problematic question items that warrant their attention for improvement. The plot shown in Figure 5 presents 12 question items whose coefficients are lower than 0.2; those 12 question numbers are displayed in the RStudio console and exported to a text file for future reference: “As seen in the discrimination plot, the following question items present a discrimination index lower than 0.2: “Q1” “Q2” “Q3” “Q4” “Q5” “Q6” “Q7” “Q8” “Q10” “Q14” “Q15” “Q17”.” Based on this information, the research team can instantly identify and discard question items Q1, Q2, Q5, Q7, and Q17, whose scores are negative, and rephrase other question items, whose scores lie between zero and 0.2, to improve the item discrimination.

In this particular study, the research team wanted to know the efficacy of computational modeling and simulations in improving students’ performance (i.e., understanding biological processes). They administered a test (with the same question items) with two groups (i.e., intervention and control) and two times (i.e., pre-test before implementing the intervention and post-test after implementing the intervention). Data preparation for analysis can be a daunting task, especially when multiple datasets are involved just as in this case study. In addition to cleaning individual datasets, the preparation entails merging and binding those multiple datasets, the processing of which takes a significant amount of time and is also vulnerable to errors.

### 4.2. One-way ANCOVA: “one_way_ancova()”

The research question about the efficacy of interventions (e.g., modeling and simulation in this case study) can be answered with the one-way ANCOVA; the function “one_way_ancova()” will conduct a one-way analysis of covariance (ANCOVA). This function automatically merges pre-post data sets, binds treatment-control data sets, runs scripts to check assumptions of one-way ANCOVA, runs the main One-way ANCOVA, and then displays and exports all outputs all at once.

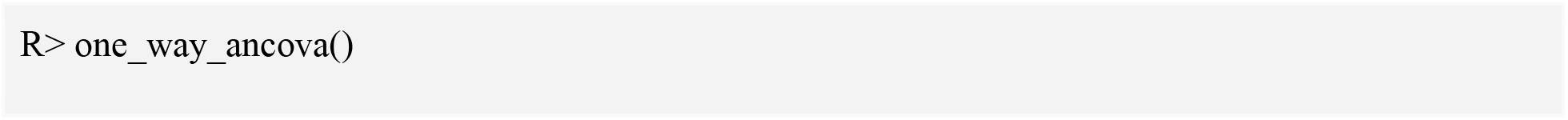

The function automatically conducts data cleaning for all four data files to be used in the one-way ANCOVA with minimal interaction with users described in the example of item analysis earlier. The outputs/results from this function are as follows. This function also automatically deletes students who skipped too many questions in all three data files. The threshold for skipped questions is interactively defined by users, as described in the previous section of item analysis.

The first output automatically displayed in the RStudio console is descriptive statistics for users to make sense of the data for analysis and use statistics in interpreting the results from this function later (Figure 6).

**Figure 6:**
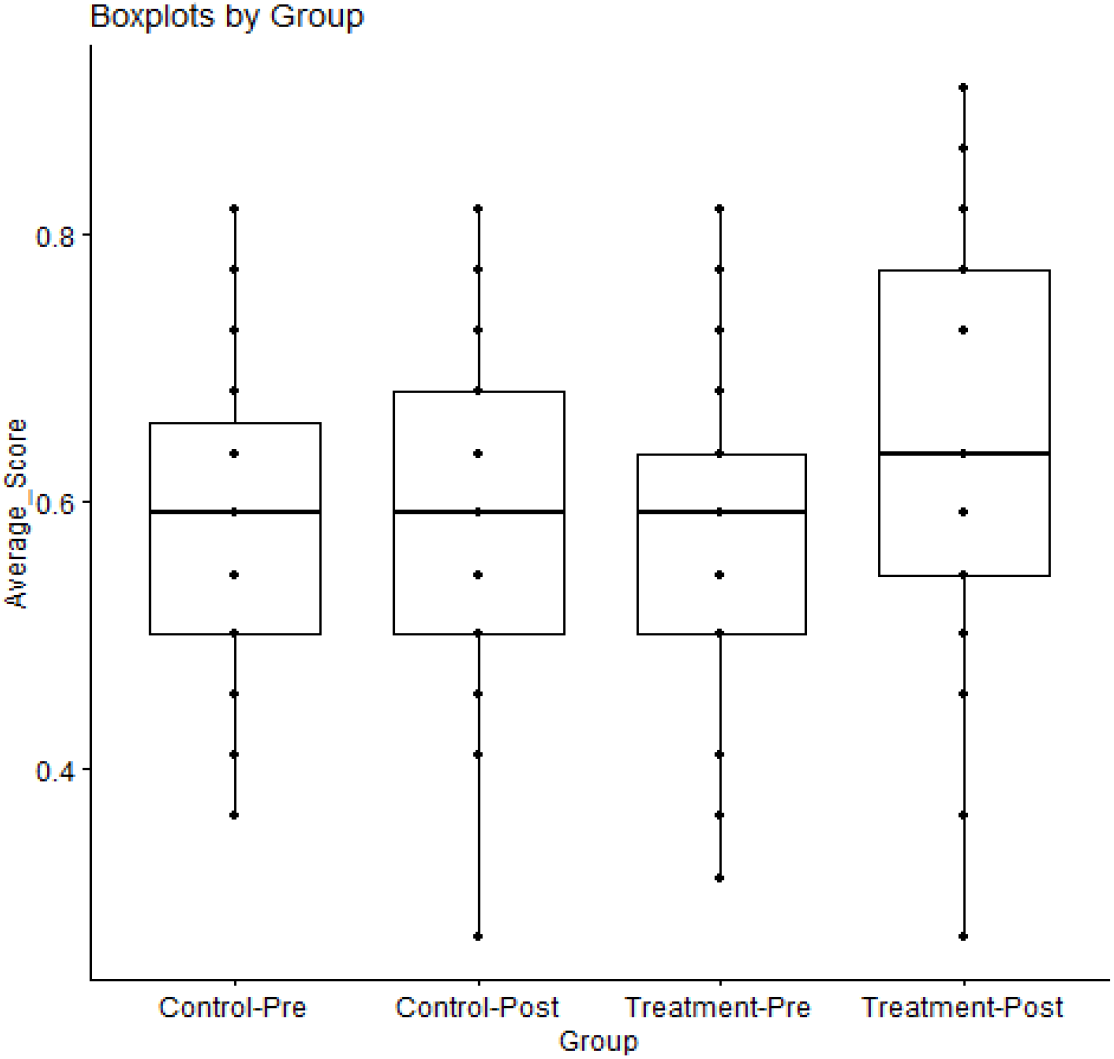
One-way ANCOVA - Boxplots

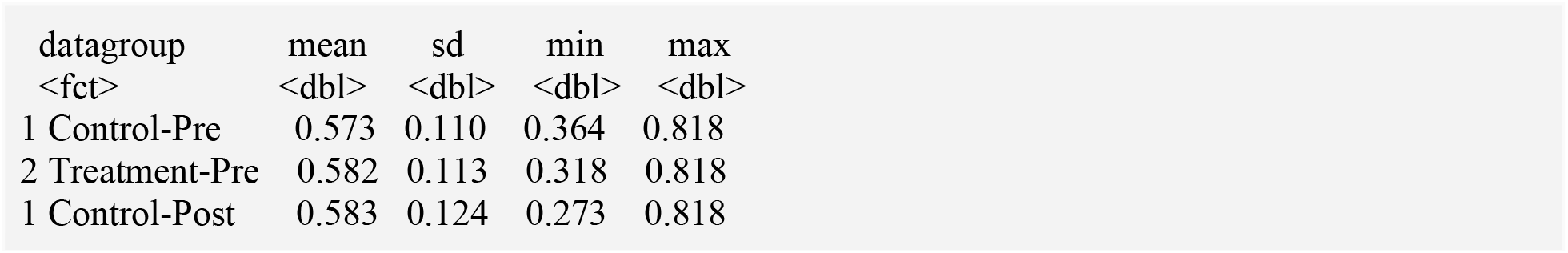

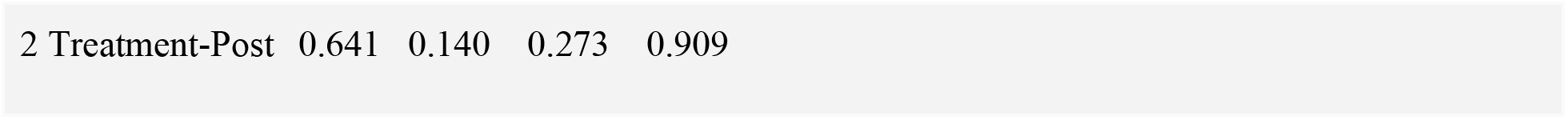

Then, the function proceeds to assumption testing. To confidently interpret the results of one-way ANCOVA, users need to check assumptions (e.g., normality of residuals, homogeneity of variance, no outlier, and homogeneity of regression line slopes) required for the parametric one-way ANCOVA. This function runs all necessary tests to generate the information for users to examine how the assumptions are satisfied and displays and exports the results as shown below.

The first assumption to check is the normality of residuals. The result of the Shapiro-Wilk normality test is presented with an interpretation in the R console (also exported to a text file in the working directory for users’ reference later).

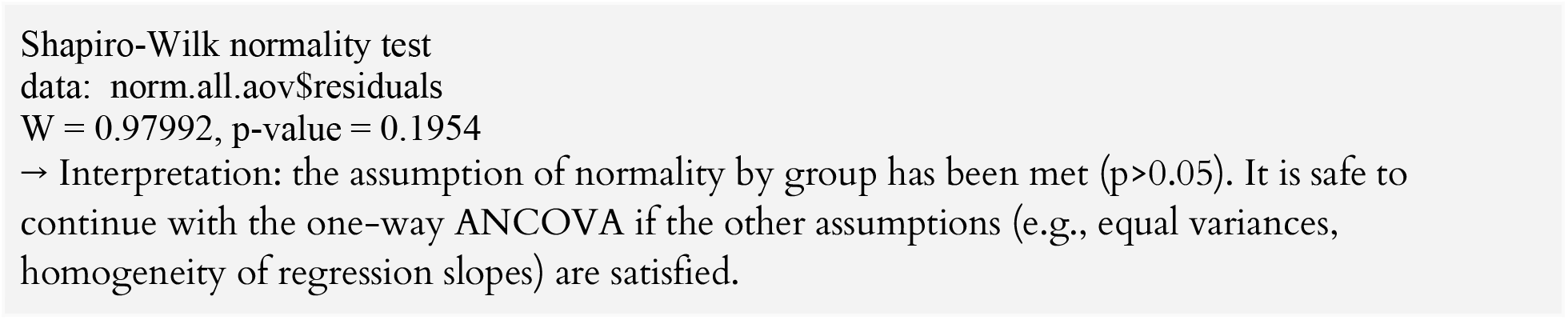

Then, the function displays a histogram and a normal Q-Q plot in the ‘Plots’ panel (also exports them as image files to the working directory) for users to examine the normality of residuals visually. A message “Refer to the histogram and the normal Q-Q plot in the ‘Plots’ panel to visually inspect the normality of residuals” is presented in the R console to direct users’ attention to the ‘Plots’ panel. Users need to refer to the normal Q-Q plot, especially if the sample size is greater than 50 because the Shapiro-Wilk test above becomes very sensitive to a minor deviation from normality at larger sample size. The function automatically displays a normal Q-Q plot in the “Plots” panel and exports it as an image to the working directory for users’ reference later. Users should see if all the points fall in the plots (refer to Figure 7 below) approximately along the reference line to assume normality confidently.

**Figure 7:**
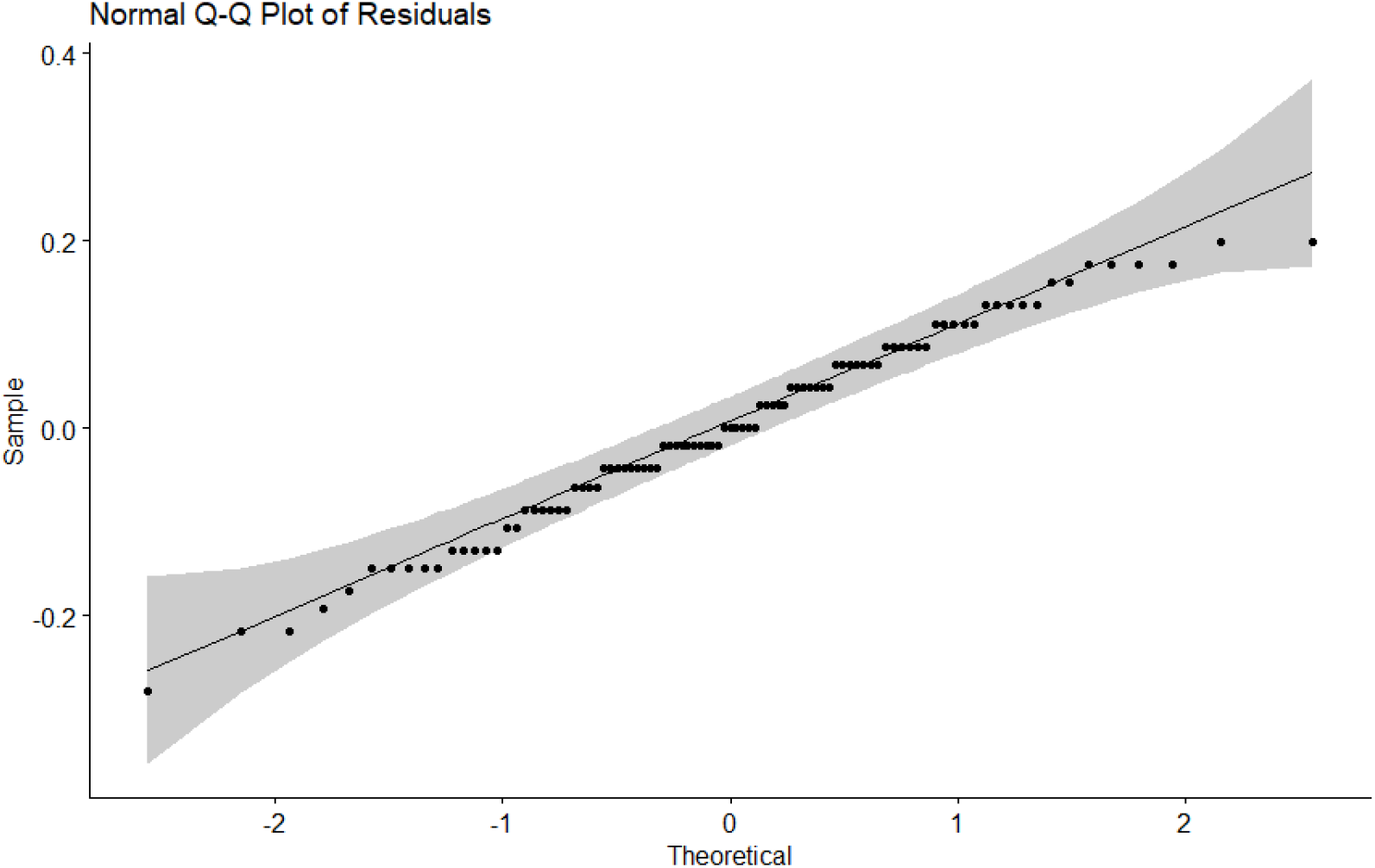
One-way ANCOVA - Normal Q-Q Plot of Residuals

The next assumption checked is the homogeneity of variance. The function runs Levene Test and reports its result with an interpretation, as shown below:

|  | Df | F value | Pr(>F) |
| --- | --- | --- | --- |
| group | 1 | 0.2977 | 0.5868 |
|  | 85 |  |  |

→ Interpretation: the assumption of equality of error variances has been met ( $p > 0.05$ ). It is safe to continue with the one-way ANCOVA if the other assumptions (e.g., normality, homogeneity of regression slopes) are satisfied.

Next, the assumption of homogeneity of regression line slopes is checked, and its result is presented with an interpretation, as shown below. The output below presents information for checking the assumption of homogeneity of regression line slopes. The output below demonstrates the sample data satisfies this assumption since the interaction term (i.e., datagroup:c_gpa) between the covariate (c_gpa) and group variable (datagroup) is not significant (p>0.05).

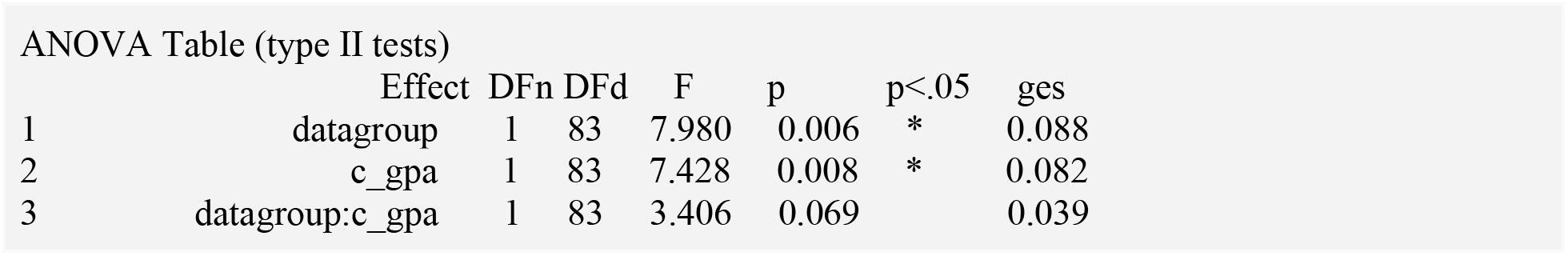

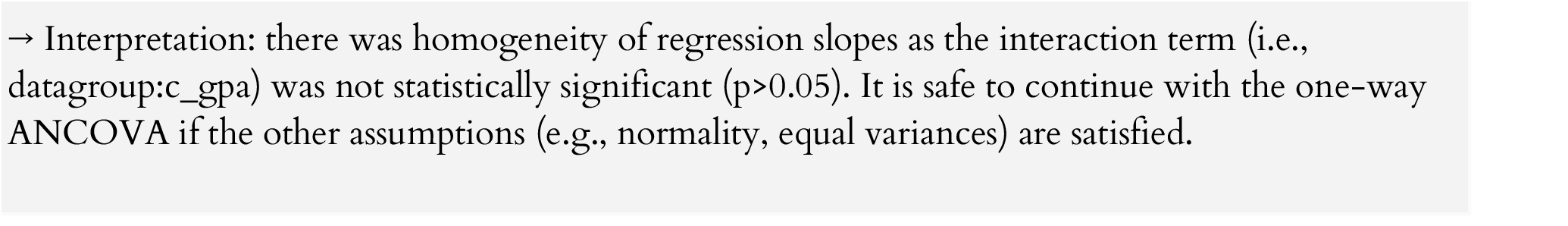

Once all assumption checks are completed, the function proceeds to run scripts to generate the main ANCOVA results. The function generates the ANOVA table (type II tests) for users to examine the ANOVA Table (type II tests). The table shows the variables of gender, native English speaker, parents’ college education, the extent of education self-funding, and pre-test scores entered as control variables in the ANCOVA model.

|  | Effect | DFn | DFd | F | p | p<.05 | ges |
| --- | --- | --- | --- | --- | --- | --- | --- |
| 1 | datagroup | 1 | 79 | 8.148 | 0.006 | * | 0.094 |
| 2 | c_gpa | 1 | 79 | 6.461 | 0.013 | * | 0.076 |
| 3 | gender | 1 | 79 | 1.584 | 0.212 |  | 0.020 |
| 4 | native_e | 1 | 79 | 0.891 | 0.348 |  | 0.011 |
| 5 | parent_c | 1 | 79 | 0.067 | 0.796 |  | 0.0009 |
| 6 | self_funding | 1 | 79 | 0.019 | 0.891 |  | 0.0002 |
| 7 | avg_score_pre | 1 | 79 | 9.775 | 0.002 | * | 0.110 |

Then, the function runs a posthoc analysis to generate estimated (or adjusted) marginal means to compare between the groups and exports a summary statement of the outputs (see the results below). The results are followed by a brief interpretation of all results so far.

|  | avg_score_pre | datagroup | emmean | se | df | conf.low | conf.high |
| --- | --- | --- | --- | --- | --- | --- | --- |
| 1 | 0.573 | Control | 0.573 | 0.019 | 84 | 0.536 | 0.611 |
| 2 | 0.582 | Treatment | 0.653 | 0.206 | 84 | 0.612 | 0.693 |

→ A sample summary: The difference of post-test scores between the treatment and control groups turned out to be significant with pre-test scores being controlled: $F(1,79)=8.148$ , $p=0.006$ (effect size=0.094). The adjusted marginal mean of post-test scores of the treatment group (0.65, SE=0.02) was significantly different from that of the control group (0.57, SE=0.02).

The ideal design to examine a pure effect of an intervention is one-way ANCOVA, which entails both pre- and post-tests and both intervention and control groups. However, it is often difficult to have all four data sets (i.e., pre- and post-test, intervention and control data). Users can conduct a paired samples t-test (or its non-parametric equivalent ‘Wilcoxon signed-rank test’ if data requirements for the parametric t-test are not satisfied) when there is only pre- and post-test data without a control group, or an independent samples t-test (or its non-parametric equivalent ‘Mann-Whitney U test’ data requirements for a parametric t-test are not satisfied) when there are only intervention and control groups without a pre-test. All these data analyses for paired samples or independent samples can be conducted with DBERlibR.

DBERlibR can also help examine the significance of changes over time; for example, a change over time can be analyzed by collecting data before (i.e., pre-test at Time 1) and after (i.e., post-test at Time 2) Intervention 1 and after Intervention 2 (i.e., post2-test at Time 3). From the context of the case study in this illustration, Intervention 1 and 2 would be modeling and simulation, respectively, enabling the research to answer another question “Will simulations further enhance students’ understanding of complex biological processes?” Data need to be collected before (Time 1) and after having students run modeling (Time 2) and after having students run simulations (Time 3), and the one-way repeated measures ANOVA can be employed to analyze data. The analysis can be time-consuming and error-prone if manually conducted. DBERlibR has this function of the one-way repeated measures ANOVA (parametric) or Freedman test (non-parametric) automatically depending on the data characteristics, as illustrated in the next section.

### 4.3. One-way Repeated Measures ANOVA: “one_way_repeated_anova()”

The function “one_way_repeated_anova()” conducts the one-way analysis of variance (ANOVA) with repeated measures at three different time points (pre-test, post-test, and post2-test). This function automatically merges pre, post, and post2 datasets, cleans data, checks all required assumptions, runs the analysis, and then displays/exports outputs for users all at once.

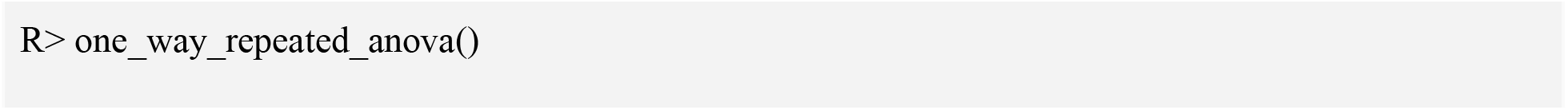

This function automatically deletes students who skipped too many questions in all three data files. The threshold for skipped questions is interactively defined by users, as described in Section 4.1.

The first output automatically displayed to the users in the RStudio console is the descriptive statistics providing an overview of the data. These statistics are represented in a box plot as well.

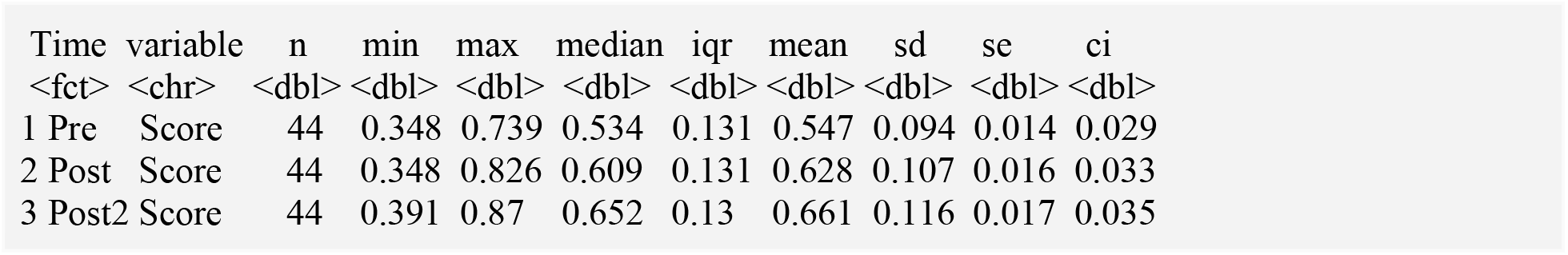

To confidently interpret the results of one-way repeated measures ANOVA, users need to check that the assumptions (no extreme outlier, normality, and sphericity) required for this parametric statistical technique are satisfied. The function runs all necessary tests to generate the needed information.

The first automatically checked assumption is the absence of outliers. The result is displayed in the RStudio console and exported in a text file for future reference. An interpretation is appended to help users understand the result, as shown below.

|  | <fct> | <fct> | <dbl> | <lgl> | <lgl> |
| --- | --- | --- | --- | --- | --- |
|  | Time | id | Score | is.outlier | is.extreme |
| 1 | Post | 44 | 0.348 | TRUE | FALSE |
| 2 | Post2 | 44 | 0.391 | TRUE | FALSE |

→ Interpretation: No extreme outlier was identified in your dataset. It is safe to continue with the one-way repeated measures ANOVA if the other assumptions (e.g., normality, equal variances) are satisfied.

The second assumption checked is the normality of residuals. The function automatically runs the Shapiro-Wilk normality test in the back end. The result is displayed in the RStudio console and exported into a text file for users’ future reference. An interpretation of the result is appended to help users understand the result, as shown below.

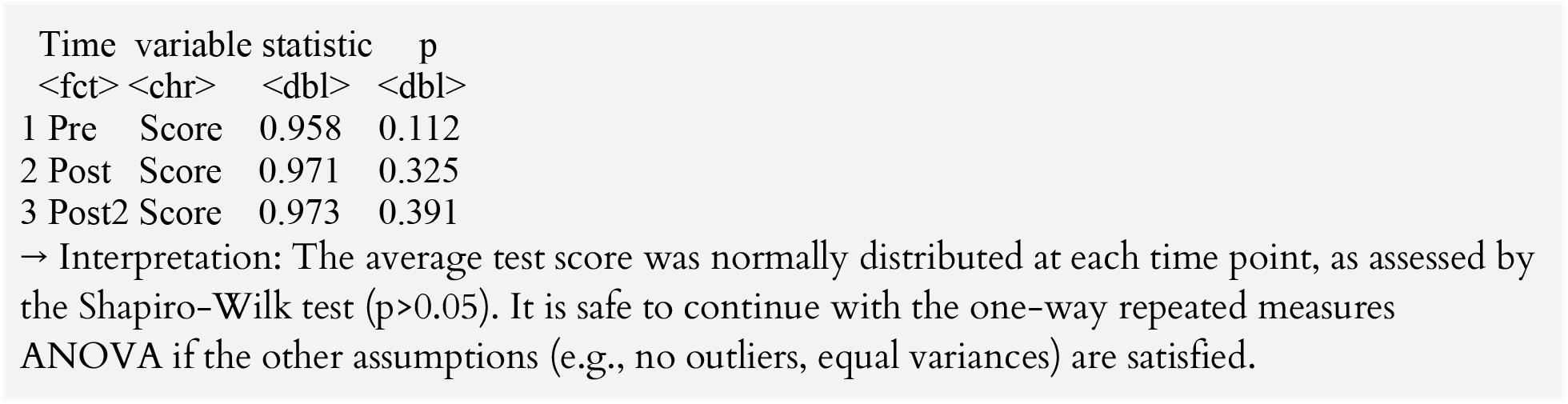

As in the other normality check, users need to visually inspect the normal Q-Q plot, especially if the sample size exceeds 50 because the Shapiro-Wilk test becomes overly sensitive to a minor deviation from normality at larger sample sizes. The function automatically displays a normal Q-Q plot in the “Plots” panel and exports it as an image to the working directory for users’ future reference. Users should ensure that all the points fall approximately along the reference line within the grayed area to assume normality confidently.

The assumption of sphericity is automatically checked during the computation of the ANOVA test (the Mauchly’s test (refer to Moulton for details) is internally run to assess the sphericity assumption). Then, the Greenhouse-Geisser sphericity correction (refer to Abdi 2010 for details) is automatically applied to factors violating the sphericity of the assumption so that users do not need to worry about this assumption.

After checking all required assumptions, the main one-way repeated measures ANOVA is performed, and its result is displayed in the RStudio console and exported into a text file for users’ future reference with an appended interpretation, as shown below. Users can safely use the result if all the required assumptions are satisfied.

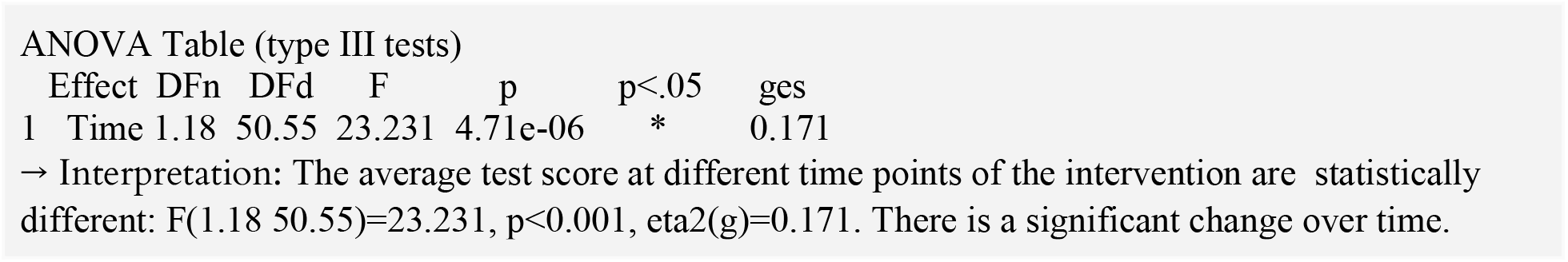

The pairwise comparisons then inform users of which intervention (time point) is more effective. An automatically generated interpretation for each pairwise comparison is appended to help users understand the result.

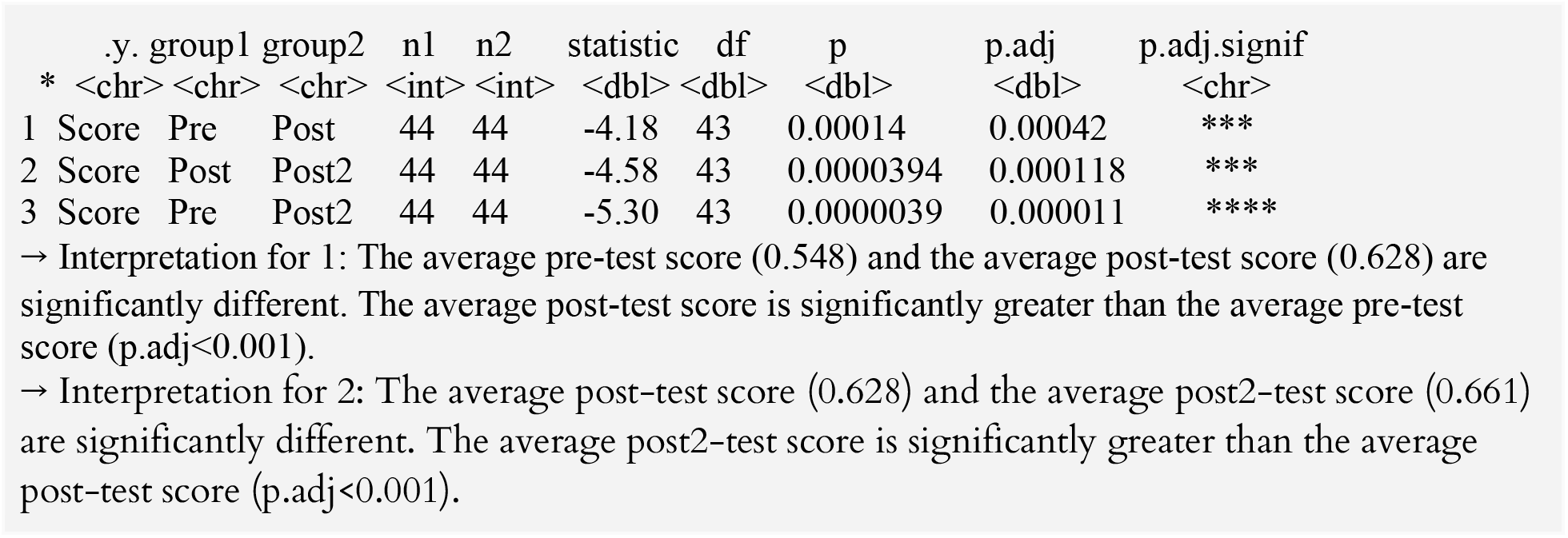

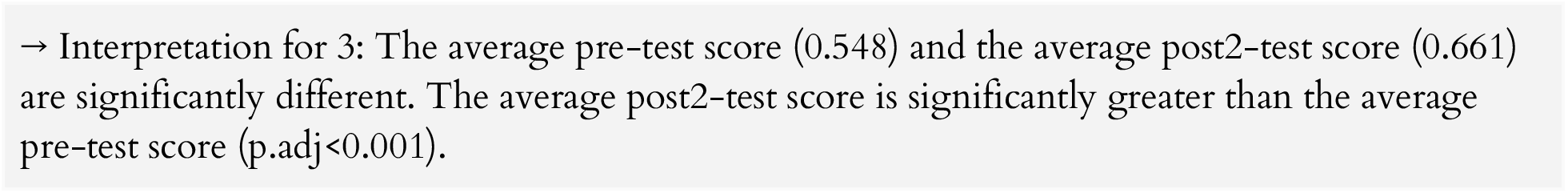

The one-way repeated measures ANOVA is known to be robust to a violation of the assumption of normality. However, this does not necessarily mean that this assumption can be overlooked since a relatively more reliable estimate(s) should be reported if available. Therefore, if the assumption of normality is violated, the function automatically proceeds to run the Friedman test, a non-parametric version of the one-way repeated measures ANOVA that can be used when the data violates the normality assumption.

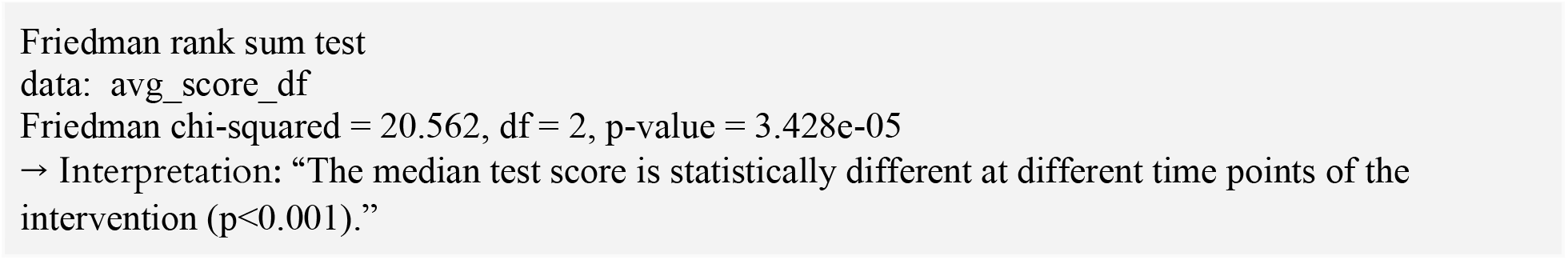

If the Friedman chi-squared is significant (i.e., p<0.05), then the function conducts pairwise comparisons.The result is displayed in the Rstudio console and exported into a text file in the working directory for users’ future reference), as shown below.

| .y. | group1 | group2 | n1 | n2 | statistic | p | p.adj | p.adj.signif |
| --- | --- | --- | --- | --- | --- | --- | --- | --- |
| * <chr> | <chr> | <chr> | <int> | <int> | <dbl> | <dbl> | <dbl> | <chr> |
| 1 Score | Pre | Post | 44 | 44 | 174. | 0.00054 | 0.002 | ** |
| 2 Score | Pre | Post2 | 44 | 44 | 108 | 0.000017 | 0.0000516 | **** |
| 3 Score | Post | Post2 | 44 | 44 | 0 | 0.000137 | 0.000411 | *** |

→ Interpretation for 1: The median pre-test score (0.544) and the median post-test score (0.609) are significantly different. The median post-test score is significantly greater than the median pre-test score (p.adj<0.001).
→ Interpretation for 2: The median pre-test score (0.544) and the median post2-test score (0.652) are significantly different. The median post2-test score is significantly greater than the median pre-test score (p.adj<0.001).
→ Interpretation for 3: The median post-test score (0.609) and the median post2-test score (0.652) are significantly different. The median post2-test score is significantly greater than the median post-test score (p.adj<0.01).

As such, the function “one_way_repeated_anove()” handles data cleaning, merging, and analysis automatically with minimal input from users and then displays/exports all results and summaries/interpretations, helping users refer to accurate information for readers and save a significant amount of time as well.

## 5. Conclusion and outlook

DBERlibR has been developed to help DBER scientists save time by automating data cleaning and analysis with minimal inputs. It enhances the reliability and validity of their findings by preventing errors, checking assumptions, showing both parametric and non-parametric results, and providing a sample interpretation for their convenience. In turn, this can help save time on cleaning and analyzing data and increase overall study reproducibility. DBERlibR provides the most frequently used analytic techniques, i.e., item analysis, paired-samples t-test (and Wilcoxon signed-rank test as necessary), independent samples t-test (and Mann-Whitney U test as necessary), one-way analysis of covariance, one-way repeated measures analysis of variance (and Friedman test as necessary), and one-way analysis of variance (and Kruskal-Wallis rank-sum test as necessary). Because of the ease of assessment data analysis, DBERlibR might entice more DBER scientists to utilize advanced statistical techniques to examine the efficacy of their educational interventions on students’ performance, especially those who are not familiar with analytic techniques. DBERlibR will contribute to the advancement of DBER by facilitating the creation and dissemination of evidence-based knowledge and practices. Short-term outcomes of DBERlibR include saving time on data cleaning and analysis and preventing errors in the results. Long-term potential outcomes include improved research reproducibility.

